# SARS-CoV-2-Triggered Mast Cell Rapid Degranulation Induces Alveolar Epithelial Inflammation and Lung Injury

**DOI:** 10.1101/2021.06.24.449680

**Authors:** Meng-Li Wu, Feng-Liang Liu, Jing Sun, Xin Li, Xiao-Yan He, Hong-Yi Zheng, Rong-Juan Pei, Hao Sun, Yan-Heng Zhou, Qi-Hong Yan, Cai-Xia Wu, Ling Chen, Guo-Ying Yu, Junbiao Chang, Xia Jin, Jincun Zhao, Xin-Wen Chen, Yong-Tang Zheng, Jian-Hua Wang

## Abstract

SARS-CoV-2 infection-induced hyper-inflammation links to the acute lung injury and COVID-19 severity. Identifying the primary mediators that initiate the uncontrolled hypercytokinemia is essential for treatments. Mast cells (MCs) are strategically located at the mucosa and beneficially or detrimentally regulate immune inflammations. Here we showed that SARS-CoV-2-triggeed MC degranulation initiated alveolar epithelial inflammation and lung injury. SARS-CoV-2 challenge induced MC degranulation in ACE-2 humanized mice and rhesus macaques, and a rapid MC degranulation could be recapitulated with Spike-RBD binding to ACE2 in cells; MC degranulation alterred various signaling pathways in alveolar epithelial cells, particularly, led to the production of pro-inflammatory factors and consequential disruption of tight junctions. Importantly, the administration of clinical MC stabilizers for blocking degranulation dampened SARS-CoV-2-induced production of pro-inflammatory factors and prevented lung injury. These findings uncover a novel mechanism for SARS-CoV-2 initiating lung inflammation, and suggest an off-label use of MC stabilizer as immunomodulators for COVID-19 treatments.

**Graphical abstract:** 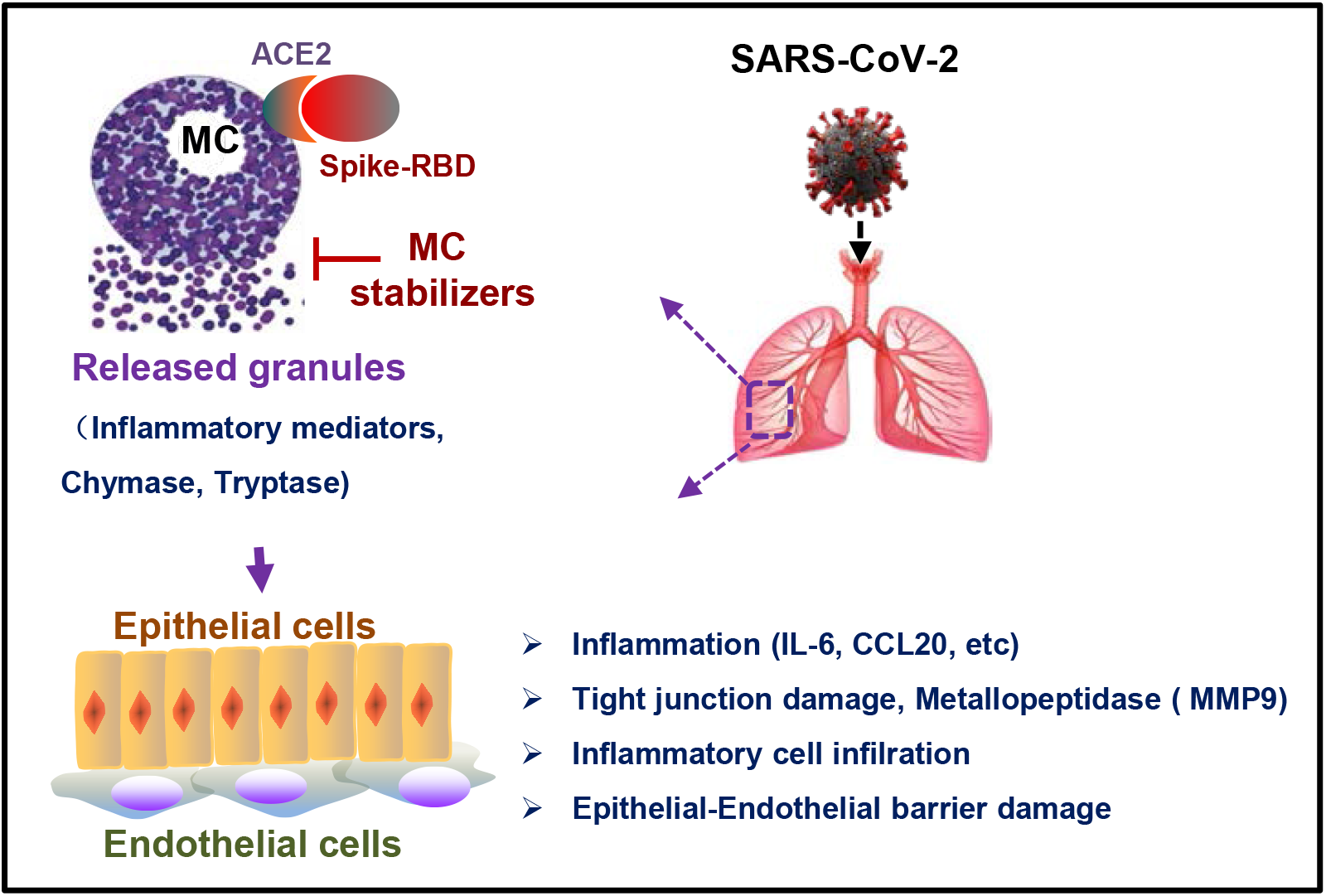

**In Brief:** SARS-CoV-2 triggers an immediate mast cell (MC) degranulation, which initiates the alveolar epithelial inflammation and disrupts the tight junction. MC stabilizers that block degranulation reduce virus-induced lung inflammation and injury.

**Highlights:**

- The binding of RBD of Spike protein of SARS-CoV-2-to ACE2 receptor protein triggers an immediate MC degranulation
- MC degranulation induces transcriptomic changes include an upregulated inflammatory signaling and a downregulated cell-junction signaling
- MC degranulation leads to alveolar epithelial inflammation and disruption of tight junctions
- MC stabilizer that inhibits degranulation reduces SARS-CoV-2-induced lung inflammation and injury in vivo

## Introduction

The Coronavirus disease 2019 (COVID-19) caused by severe acute respiratory syndrome coronavirus 2 (SARS-CoV-2) has become a global pandemic. The common clinical symptoms of SARS-CoV-2 infection include fever, cough, shortness of breath and dyspnea, and severe cases rapidly progress to acute respiratory distress syndrome, multiple organs failure and death ^1–4^. A higher incidence of severity and deaths in older individuals has been observed ^5^. Currently, no specific therapeutic agents are approved for the COVID-19 ^6,7^; several preventive vaccines have been granted emergency use by WHO and deployed globally.

A key pathologic feature of COVID-19 is the hyper-inflammatory response (also termed hypercytokinemia or ‘‘cytokine storm”) in association with severe COVID-19 disease, and those inflammatory cytokines and chemokines produced in *vivo* lead to damage to the alveoli epithelial cells and capillary endothelial cells ^8–16^. Known inflammatory cytokines and chemokines that produced in excess in patients with severe diseases include IL-6, IL-8, IL-1β, TNF-α, IFN-γ, MIP1α and 1β, CCL2, CCL5, CCL20, CXCL1, CXCL2, CXCL8, CXCL10 and CXCL17 ^10,14–21^. Among them, elevated IL-6, TNF-α and C-reaction protein (CRP) levels have been shown to be independent risk factors for the severity of COVID-19 disease ^14,18,21–23^.

Therefore, targeting the hyper-inflammation loop to develop the immunomodulatory agents is an attractive strategy for developing treatment for severe COVID-19 cases ^8,24,25^. Some drug candidates have been tried in order to mitigate hyper-inflammation and improve clinical outcomes. These include humanized anti-IL-6 receptor antibody Tocilizumab ^26^, anti-IFN-γ monoclonal antibody Emapalumab, IL-1 receptor antagonist Anakinra (NCT04324021), JAK1 and JAK2 inhibitors Baricite and Rituxolitinib ^27^ (NCT04338958), and the Corticosteroids such as Dexamethasone ^28–30^. But none of which worked well thus far, Tocilizumab was not effective for preventing intubation or death in moderately ill hospitalized patients ^31,32^; another IL-6 receptor monoclonal antibodies Sarilumab not only failed to improve clinic outcomes and reduce mortality, but led to serious complications ^33^. Of note, despite the probable benefit of calming the cytokine storm, to suppress inflammation and immune response may impede on viral clearance ^34^. Indeed, corticosteroid treatment suppresses the overall immune responses, but also impairs the induction of anti-viral type-I interferon responses ^35,36^. Therefore, the development of effective immunomodulatory agents that suppress inflammation, without compromising host immune protection, is required.

Lung is the primary target for the corona viruses. The alveoli of the lung are covered with angiotensin-converting enzyme 2 (ACE2) - expressing epithelial cells. It is reasonable to hypothesis that the alveoli epithelial inflammation is a prerequisite for damage of both alveoli epithelial cells and capillary endothelial cells during SARS-CoV-2 infection ^37–39^, during which infected epithelial cells recruit and activate monocytes and macrophages to secrete pro-inflammatory cytokines, which further recruits neutrophils and activates T-lymphocytes to exacerbate inflammation ^13^. As shown in the postmortem lung biopsies, there were diffuses alveolar damage with necrosis of alveolar lining cells, pneumocyte type 2 hyperplasia, and linear intra-alveolar fibrin deposition, widespread vascular thrombosis with microangiopathy and occlusion of alveolar capillaries ^40,41^. Therefore, uncovering the mechanisms of epithelial inflammation caused SARS-CoV-2 is of uppermost priority.

Mast cells (MCs) are tissue resident cells that strategically placed throughout the host-environment interface including the whole respiratory tract and the nasal cavity. Besides being as the main effector cells in allergy, MCs can interact with various immune cells through release of soluble factors or direct contact, and link innate and adaptive immunity ^42–44^. In SARS-CoV-2 infection, the postmortem lung biopsies of COVID-19 patients show a massively increased density of perivascular and septal MCs, suggesting MCs were recruited to the alveolar septa to play certain unknown functions ^45,46^.

To investigate the role of MCs in SARS-CoV-2 infection, we examined MC degranulation-induced alveolar epithelial inflammation and lung injury. By using rhesus macaques and ACE-2 humanized mice, we found that SARS-CoV-2 infection induced MC degranulation in the peri-bronchus. Further analysis in cell cultures revealed that the binding of Spike-RBD to ACE2 triggered an immediate MC degranulation (as short as 5 min), which led to the production of pro-inflammatory factors and the disruption of tight junctions of alveolar epithelial cells. Importantly, the administration of clinical MC stabilizers that block degranulation reduced SARS-CoV-2-induced production of pro-inflammatory factors and prevented lung injury. These findings uncovers a novel mechanism of SARS-CoV-2 infection initiated alveolar epithelial inflammation and inducing lung Injury, and suggests an off-label use of MC stabilizers as the immunomodulators to treat COVID-19.

## Results

### SARS-CoV-2 induces MC degranulation in lung of ACE2-humanized mice

To investigate whether SARS-CoV-2 can induce MC degranulation in *vivo*, the ACE2-humanized inbred mice^47^, termed C57BL/6N-Ace2^em2(hACE2-WPRE,pgk-puro)/CCLA^, were intranasally infected with SARS-CoV-2 (strain 107) at a dose of 2×10^6^ TCID_50_, then euthanized at the different days post-infection (dpi) to harvest lung tissues for histological analysis. Our results confirmed previous observation that SARS-CoV-2 virions were mainly distributed in the peri-bronchus and bronchioalveolar-duct junction ^47–49^, as shown by immunostaining of SARS-CoV-2 nucleocapsid protein at 1- and 3-dpi (Figure S1). MCs and their degranulation are indicated by the metachromatic labeling with Toluidine blue (T. blue) ^50^. Compared with mock-infection (Figure 1A), at the 1 dpi, the accumulation of MCs in peri-bronchus was observed and the release of granules was found in the alveolar space (Figure1C). The progressive MC degranulation over the course of SARS-CoV-2 infection could be seen with more abundance of released granules distributed more widely in alveolar spaces (Figure1C, 1E and IG). The lung lesions around the area of MC accumulation and degranulation were examined by staining with Hematoxylin and eosin (H.E.). Compared with mock-infection control (Figure 1B), lung lesions, including inflammatory cells (lymphocytes and monocytes) infiltration, hemorrhage, alveolar septal thickening, and mucosa desquamation, were observed around the areas of MC accumulation and degranulation, and appeared progressively worsening over the course of SARS-CoV-2 infection (Figure 1D, 1F and 1H).

**Figure 1.**
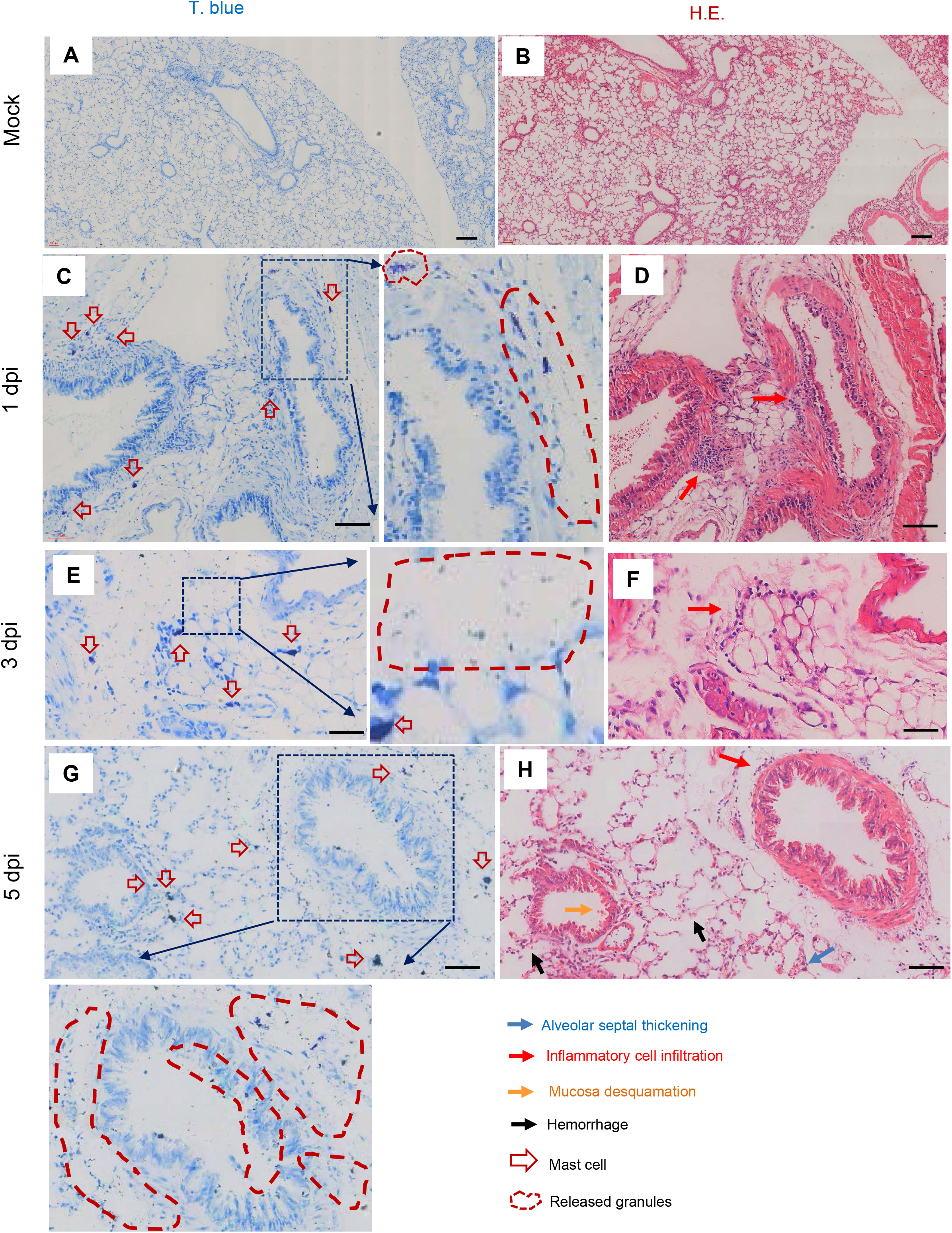
SARS-CoV-2 induced mast cell degranulation and lung injury in hACE2-humanized mice. Five C57BL/6N-Ace2^em2(hACE2-WPRE,pgk-puro)/CCLA^ mice were intranasally infected with SARS-CoV-2 (strain 107) at a dose of 2×10^6^ TCID_50_, two mice were used as the mock-infections. The mice were euthanized at the 1 dpi, 3 dpi and 5 dpi, and the lung tissues were harvested for histological analysis. Toluidine blue staining was used to observe MCs and their degranulation, and the lung injury was observed by H.E. staining. Scale bar: 100 μm.

To ensure the above observations are not limited to one experimental model of SARS-CoV-2 infection, another mouse model, namely Ad5-hACE2-transduced BALB/c mice ^51^, were used. The mice were intranasal challenge with 7×10^4^ TCID_50_ SARS-CoV-2, then sacrificed at different times to harvest the lungs for histological staining. Similar results were obtained. Compared with the mock-infection (Figure 2A), SARS-CoV-2 challenge induced a progressive MC degranulation in the peri-bronchus and bronchioalveolar-duct junction (Figure 2C, 2E and 2G). Simultaneous H.E. staining of the adjacent lung section showed lung lesions around these areas (Figure 2D, 2F and 2H), but not in mock-infected mice (Figure 2B). Taken together, these data demonstrate convincingly that SARS-CoV-2 infection induces MC accumulation and degranulation around the area of lung lesion in ACE2-humanized mice.

**Figure 2.**
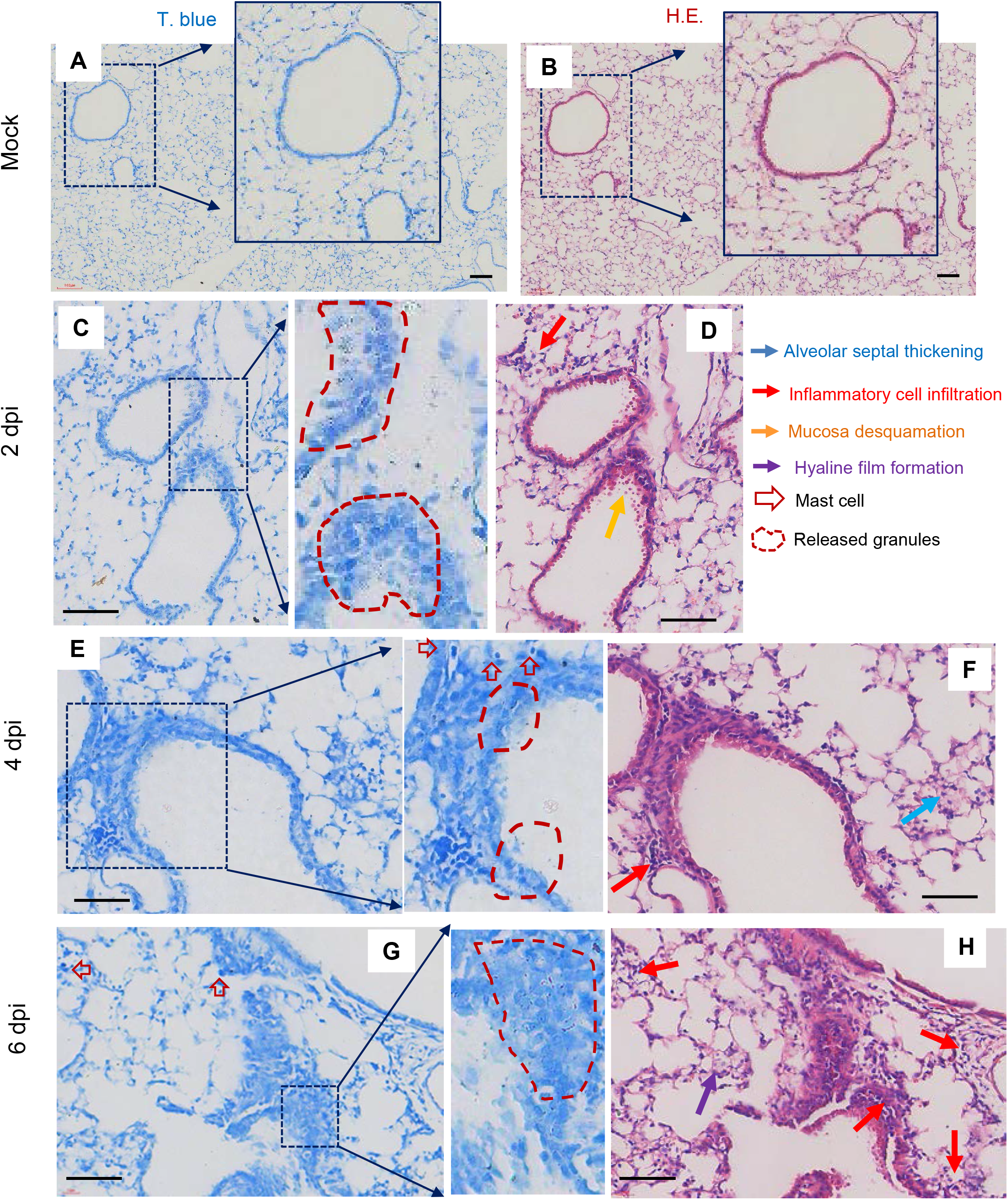
SARS-CoV-2 induced mast cell degranulation and lung injury in Ad5-hACE2-transduced BALB/c mice. Five Mice were transducted with 2.5×10^8^ FFU of Ad5-hACE2 in 75 mL of DMEM intranasally, 5 days later, mice received 7×10^4^ TCID_50_ of SARS-CoV-2. Two mice were used as the mock-infections. The lungs were collected at 2 dpi, 4 dpi, and 6 dpi. Toluidine blue staining was used to observe MCs and their degranulation, and the lung injury was observed by H.E. staining. Scale bar: 100 μm.

### SARS-CoV-2 also induces MC degranulation in lung of rhesus macaque

Having firmly established a pathogenesis model in mice, we went on the investigated whether it is producible in nonhuman primates, which have often been consider a translational gate-keeper before any scientific discovery is progressed to human testing. The SARS-CoV-2 can infect Chinese rhesus macaques (chRMs) (*Macaca mulatta*) and recapitulate much of the characteristics of immunological pathogenesis of human COVID-19; importantly, aged chRMs showed the delayed but more severe cytokine storm and higher immune cell infiltration, compared with young chRMs^52^. This model is therefore most suited for in-depth dissection of inflammatory responses in relation to MC activation.

We first confirmed that SARS-CoV-2 infection led to MC degranulation in lungs of chRMs. The experiment was performed by anesthetize young chRMs (3- to 6-year old) or aged chRMs (17- to 19-year old) first, and then infect them intratracheally with SARS-CoV-2 (1□×□10^7^ TCID_50_) (strain 107). The animals were sacrificed at 7 or 15 dpi, and lungs were harvested and sectioned for T. blue staining to examine changes in MCs. Compared with the mock-infections (Figure 3A, 3G), SARS-CoV-2 challenge recruited MCs accumulation in the peri-bronchus and bronchioalveolar-duct junction at both 7- and 15-dpi, with noted MC degranulation (Figure 3B-F; Figure 3H-3J). As expected, compared with young chRMs (Figure 3B-3F), aged chRMs have showed more SARS-CoV-2-induced MCs accumulation and degranulation (Figure 3H-3J). Taken together, we demonstrated that SARS-CoV-2 induces MC degranulation in lung of rhesus macaque, similar to that in mice.

**Figure 3.**
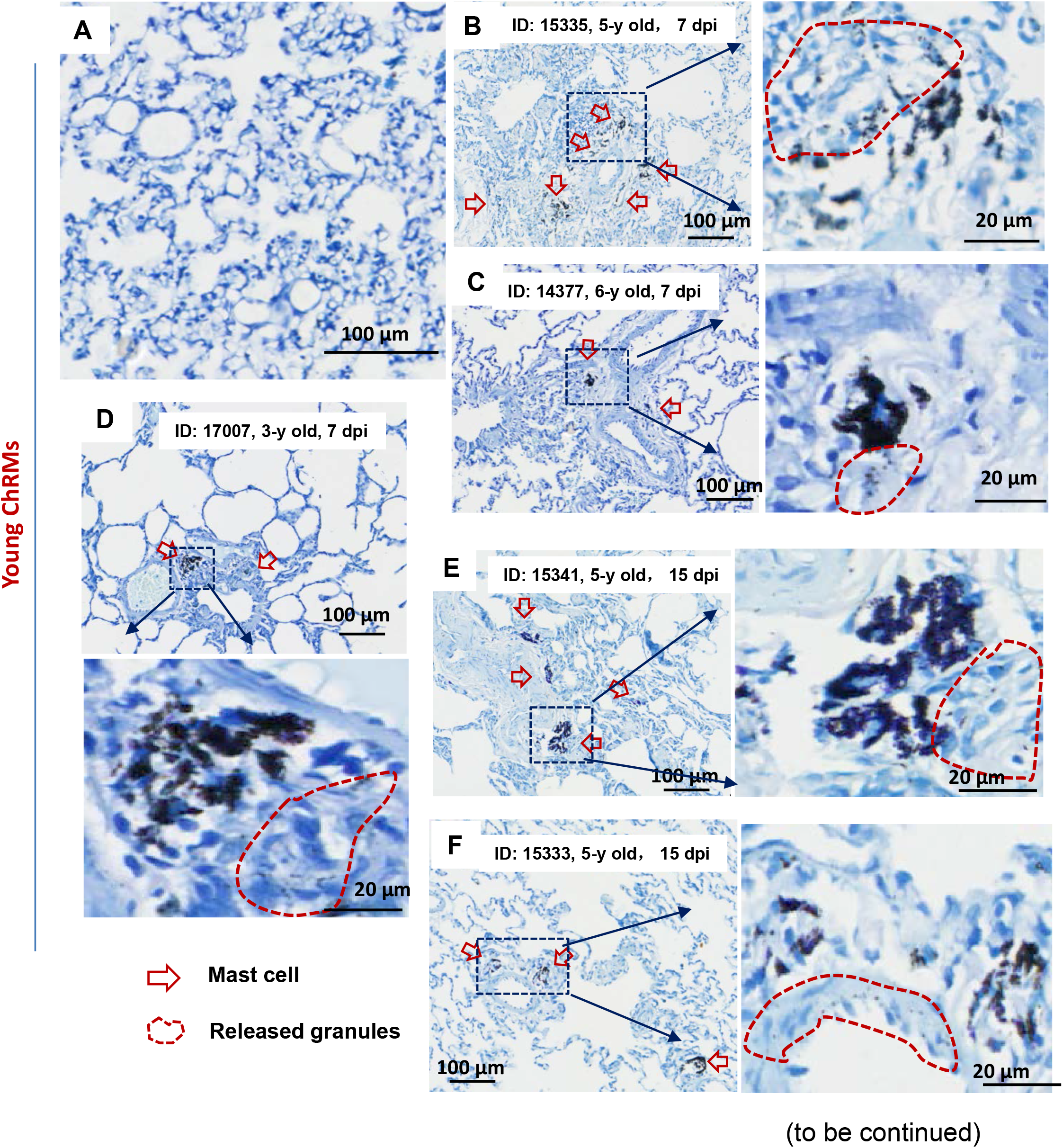
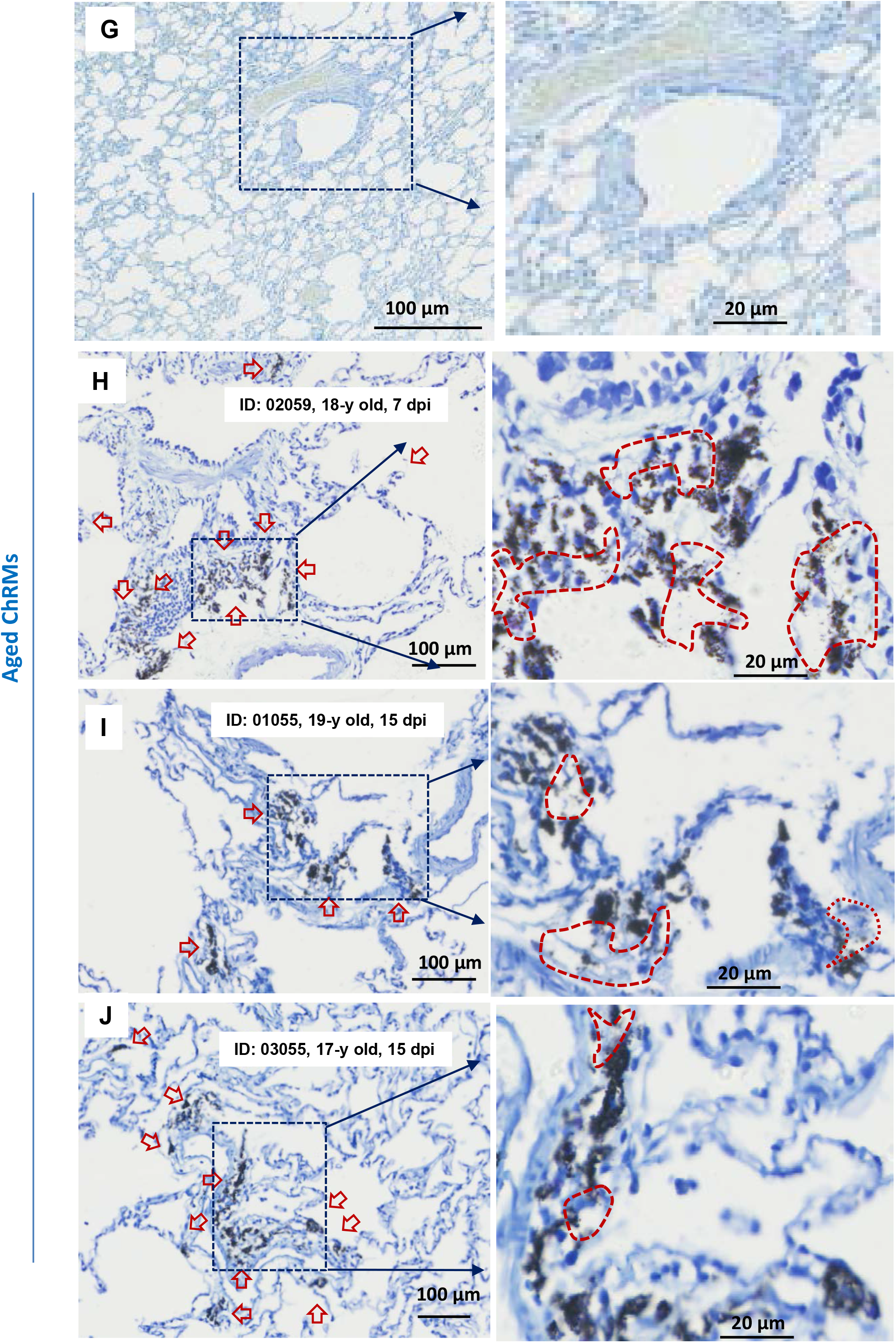
SARS-CoV-2 induced mast cell degranulation in chRMs. Eight chRMs, including young group (3- to 6-year old) and aged group (17- to 19-year old), were anaesthetized by Zoletil 50 and intratracheally inoculated with SARS-CoV-2 (1□×□10^7^ TCID_50_) in a 2 mL volume by bronchoscope. The animals were euthanized at 7 or 15 dpi and the lung lobes were collected for histology analysis. Toluidine blue staining was used to observe MCs and their degranulation.

### The binding of Spike-RBD to ACE2 triggers a rapid MC degranulation

In order to further investigate the molecular mechanism responsible for SARS-CoV-2 infection caused MC degranulation, we chose human MC cell line LAD2 cells. LAD2 cells were treated with SARS-CoV-2 (M.O.I. = 1) (strain 2019-nCoV WIV04) ^53^, and MC degranulation was assessed by the reduction of the intensity of immunostaining of cytoplasmic avidin granules as described ^50,54^. Results indicate SARS-CoV-2 induced an immediate LAD2 cell degranulation as shown by rapid decrease of immunostaining of cytoplasmic avidin granules within 5 min of virus infection, and the induction of cellular degranulation progressed over the course of 2 h viral infection (Figure 4A).

**Figure 4.**
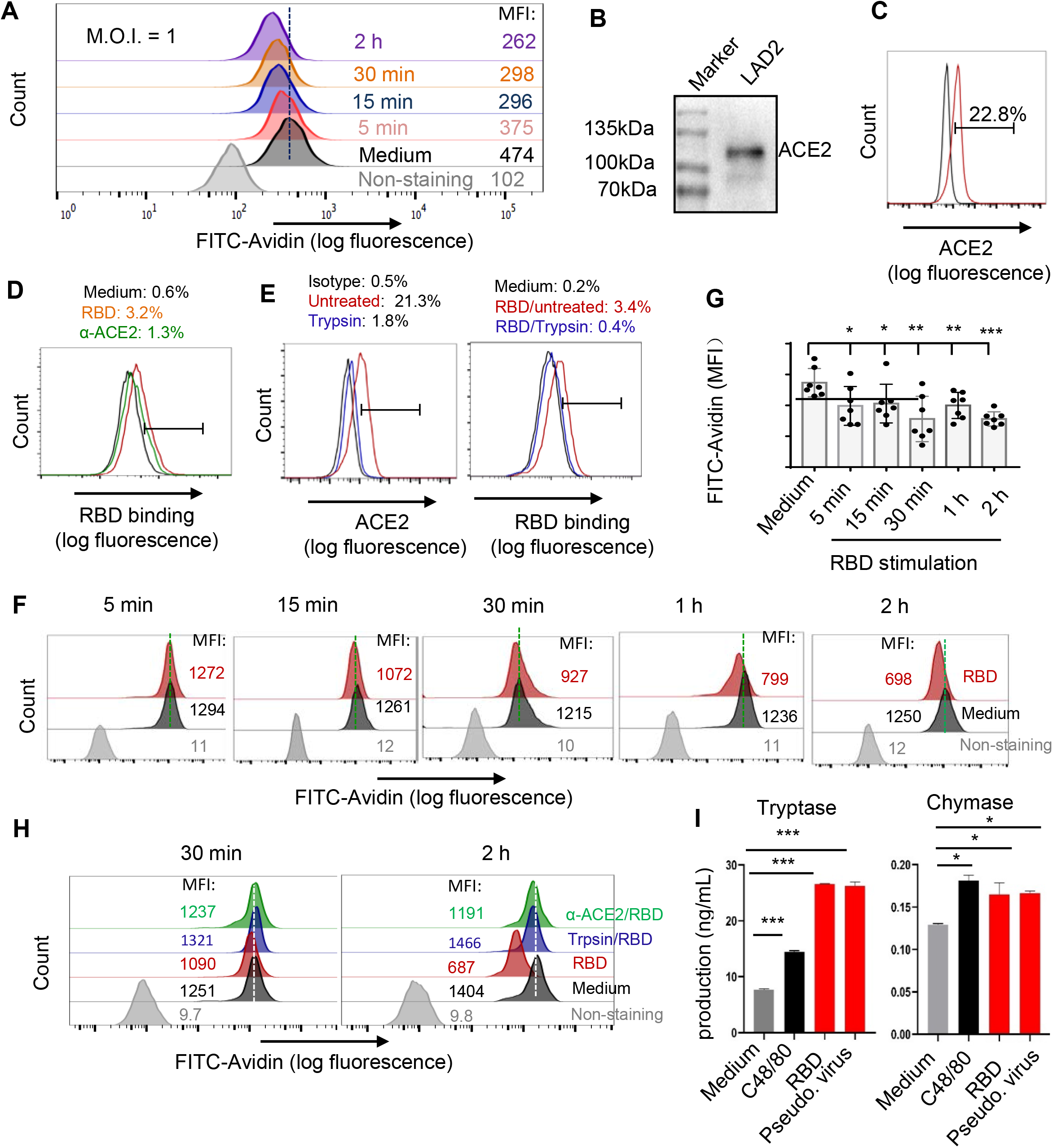
The binding of Spike-RBD to ACE2 triggers a rapid MC degranulation. (**A**) SARS-CoV-2 (M.O.I. =1)-induced LAD2 degranulation, detected by flow cytometry with immunostaining the intracellular avidin. (**B**, **C**) ACE2 expression in LAD2 cells, detected with Western blotting and flow cytometry with immunostaining with specific antibodies. (**D**, **E**) Prior-blocking with anti-ACE2 antibody or treatment with trypsin to remove ACE2 reduces RBD binding. LAD2 cells were incubated with anti-ACE2 antibody (5 μg/ml) at 37°C for 1 h (D), or LAD2 cells were treated with 0.25% trypsin (without EDTA) for 10 min at 37 °C (E), then Spike-RBD (5 μg/ml) were added for binding at 4°C for 1h, and the binding of Spike-RBD to LAD2 cells was detected with flow cytometry. (**F, G**) Spike-RBD induces LAD2 degranulation. LAD2 cells were incubated with Spike-RBD (5 μg/ml) at 37°C for the indicated time, then cells were fixed with 4% paraformaldehyde and permeabilized and immunostained with anti-avidin-FITC at 4°C for 1 h, and analyzed with flow cytometry. Results from independent repeats were summarized and presented (G). (**H**) Removing the binding of Spike-RBD losses the induction of LAD2 cell degranulation. LAD2 cells were prior-treated with anti-ACE2 antibody or trypsin as above, then Spike-RBD (5 μg/ml) was added for the indicated time, and cell degranulation was detected with flow Cytometry. (**I**) The degranulated components. LAD2 cells were treated with Spike-RBD (5 μg/ml) at 37°C for 2 h, and the supernants were harvested to detect the released Tryptase and Chymase by ELISA. Data are presented as mean ± SD. * p<0.05, ** p<0.01 and *** p<0.001 are considered significant differences. MFI: mean fluorescence intensity.

Because SARS-CoV-2 infection is mediated by Spike protein which provides first contact to ACE2 receptor on susceptible cells, we focused on this molecular interaction first. LAD2 cells express modest levels of ACE2 receptor as demonstrated by Western blotting (Figure 4B) and flow cytometry (Figure 4C) by using anti-ACE2 specific antibody, and LAD2 cells are susceptible to SARS-CoV-2 infection (Figure S2). The recombinant receptor-binding domain (RBD) of SARS-CoV-2 Spike glycoprotein can bind to LAD2 cells at 4°C, and prior-blocking with anti-ACE2 specific antibody prevents the Spike-RBD binding (Figure 4D). Likewise, the removal of surface-expressed ACE2 with trypsin treatment significantly diminished the Spike-RBD binding (Figure 4E).

Spike-RBD stimulation did not show the obvious elevation of the *de novo* synthesis of cytokines in LAD2 cells within a short 2 h stimulation (Figure S3), whereas, Spike-RBD binding to ACE2 is accompanied by an immediate degranulation, as shown by the significant reduction of intracellular avidin intensity within 5 min of Spike-RBD treatment (Figure 4F, 4G), and the degranulation progressively increase over the course of 2 h period of incubation (Figure 4F, 4G). The nucleocapsid protein treatment did not induce LAD2 degranulation (Figure S4). The above results indicate that the observed rapid LAD2 cell degranulation is triggered by Spike-RBD binding to ACE2 receptor protein.

To confirm the specific interaction between Spike-RBD and ACE2 is key to MC degranulation, we blocked the interaction by either prior-treatment with anti-ACE2 antibody or trypsin treatment to remove surface-expressed ACE2 and found Spike-RBD treatment of LAD2 cells no longer able to induce degranulation (Figure 4H).

Spike-RBD-induced LAD2 degranulation was also quantified by the release of intracellular Tryptase and Chymase. The treatments of LAD2 cells with either Spike-RBD or Spike-pseudotyped lentivirus (Pseudo. Virus) for 2 h significantly increased the release of Tryptase and Chymase, to the levels similar or even higher than that treated with the compound 48/80 (C48/80) which was used as a positive experimental control for stimulation of MC degranulation (Figure 4I). Taken together, these data demonstrate that the binding of Spike-RBD to ACE2 triggers an immediate MC degranulation, which is rapid and specific.

### Transcriptome analysis reveals MC degranulation-induced a remodeling of cellular signalings in human alveolar epithelial cells

SARS-CoV-2 infection causes severe alveolar epithelial inflammation and barrier dysfunction ^55,56^, of which MC degranulation may play a role. To test whether this is true, we performed transcriptome analysis in the Spike-RBD-induced MC degranulation model. The culture supernatant of Spike-RBD-treated LAD2 cells were harvested and used to treat A549 cells (an adenocarcinomic human alveolar basal epithelial cell line) for 24 h, then the transcriptome in A549 cells were analyzed using standard protocols. Data from 3 independent repeats were summarized. The volcano plot displayed a total of 1, 667 up-regulated genes and 907 down-regulated genes after A549 cells were treated with supernatants from Spike-RBD-treated LAD2 cells (Figure 5A). Gene ontology (GO) functional enrichment analysis of differently expressed genes (DEGs) showed obvious upregulation of gene sets that regulate cytokines signaling and production, the activation of myeloid and leukocyte, cell adhesion, innate immune and inflammatory response, and cell apoptotic signaling; in contrast, the gene sets that regulate cell cycle, cell division, and the actin filament-based cell movement processes were downregulated (Figure 5B).

**Figure 5.**
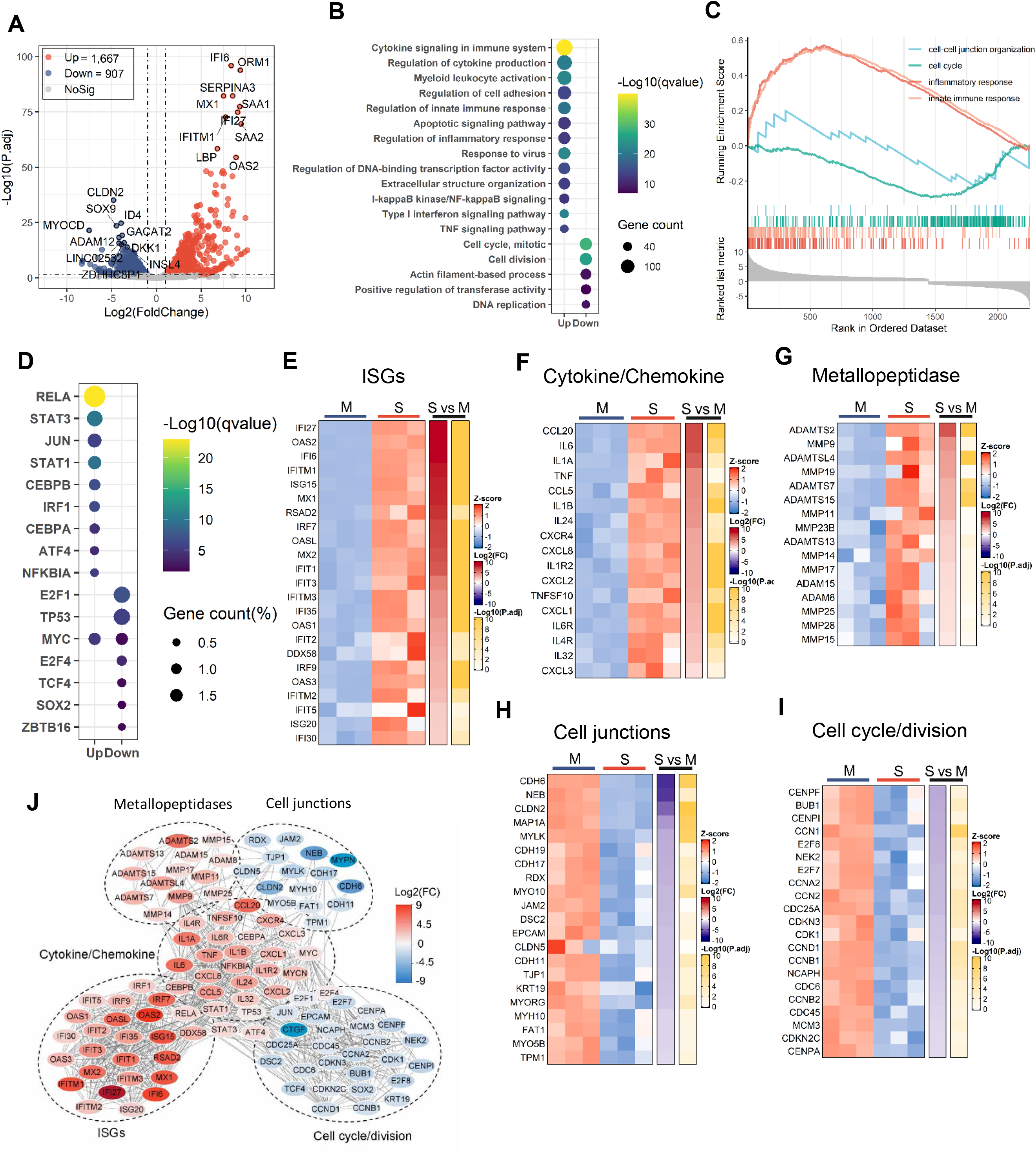
Transcriptome analysis of A549 cells treated with RBD/LAD2 supernatant. (**A**) Volcano plot of differentially expressed genes (DEGs) comparing RBD/LAD2 supernatant versus medium. The symbols of top10 up-regulated or down-regulated genes are shown. (**B**) Gene ontology (GO) functional enrichment analysis of DEGs. The color bar indicates the minus logarithm of *q* values, and bubble size indicates the absolute gene counts enriched in a GO term. (**C**) Gene set enrichment analysis (GSEA) showing the distribution of the gene sets that related to inflammatory response, innate immune response, cell-cell junction organization or cell cycle and the enrichment scores based on DEGs. (**D**) Transcription-factor enrichment analysis of DEGs. The color bar indicates the minus logarithm of *q* values, and bubble size indicates the gene enrichment ratio regulated by a transcription factor. (**E-I)** Heatmaps showing relative expression level (left panel), fold change (middle panel), and adjusted p values (right panel) for sets of ISGs (E), cytokine- and chemokine-related genes (F), Metallopeptidase (G), cell junction-related genes (H), cell cycle- and division-related genes (I). M, medium; S, RBD/LAD2 supernatant. (**J**) A Protein-Protein interaction network analysis of the core DEGs. The color bar represents the fold change of protein-coded genes at transcriptome level.

The gene set enrichment analysis (GSEA) linked the upregulated genes to the regulation of inflammatory and innate immune responses, and the downregulated genes to the regulation of cell cycles and cell-cell junctions (Figure 5C). Transcription-factor enrichment analysis of DEGs showed that the upregulated transcription factors were mainly those governing the immune and inflammatory response, e.g., *RELA*, *STAT3*, *STAT1*, *JUN*, *CEBP-*α*/*β, *IRF1*, and *ATF4;* and the down-regulated transcription factors were mainly linked to the regulations of cell proliferation and metabolisms, such as *E2F1*, *E2F4*, *TP53*, and *ZBTB16* (Figure 5D).

The core DEGs with highly differential expression were catalogized. The stimulation with RBD/LAD2 supernatants induced elevated expression of antiviral immune genes in A549 cells, and a significant upregulation of genes for ISGs and IFN-I responses (e.g., *IFI27*, *OAS2*, *IFI6*, *IFITM1*, *ISG15*, *MX1*, *RSAD2*, *IRF7*, *OASL*, *MX2*, *IFIT1*, and *IFIT3*) (Figure 5E) has been observed. Other prominent features were the upregulation of pro-inflammatory cytokines/chemokines, particularly *CCL20*, *IL-6*, *IL1*α, *TNF*, *CCL5*, *IL1*β and *CXCL8* (*IL-8*) genes (Figure 5F), and an upregulation of metallopeptidase-encoding genes (e.g., *ADAMTS2* and *MMPs*) (Figure 5G). RBD/LAD2 supernatants induced a downregulation of a large number of genes encoding regulators of cell junctions and cell cycles/division. The downregulated core DEGs are related to genes code for cell junctions included the tight junction proteins (e.g., *CLDN2*, *CLDN5*, *TJP1*, and *JAM2*), the cadherins (e.g., *CDH6*, *CDH19*, *CDH17*, *CDH11*, *FAT1*, *DSC2*, and *EPCAM*), the cytoskeleton/microtubule-associated proteins (e.g., *NEB*, *KRT19*, *RDX*, *TPM1*, and *MAP1A*), and myosins (e.g., *MYH10*, *MYO10*, *MYO5B*, *MYLK*, and *MYORG*) (Figure 5H). Among them, the downregulated core DEGs linked to cell cycles/division were mainly those encode for the mitotic Serine/Threonine kinases and their regulators (e.g., *BUB1*, *NEK2*, *CDK1; CDKN3*, *CDKN2C*), centromere proteins (e.g., *CENPF*, *CENPI*, *CENPA*), transcription factors and cyclin family members that regulate transition phase of the cell cycle (e. g., *E2F7*, *E2F8; CCNA2*, *CCND1*, *CCNB1*, *CCNB2*), genome replication regulators (e.g., *CDC45*, *CDC6*, *MCM3*), and cell division regulators (e.g., *CCN2*, *CDC25A*), *etc* (Figure 5I). A Protein-Protein interaction network (PPI) analysis of the core DEGs was constructed to visualize the relationship between DEGs and between signaling pathways (Figure 5J).

Taken together, the transcriptome data reveal that MC degranulation significantly alters many cellular signaling pathways in human alveolar epithelial cells, particularly those that upregulate immune responses and inflammation and those that downregulate signaling related to cell junction proteins and cell division.

### Inhibition of MC degranulation abolishes alveolar epithelial inflammation and protects the tight junction proteins

To sought drug candidates that may modulate MC degranulation and consequential cytokine release, we examined known MC stabilizers including the second generation antihistamines Ebastine (Eba) and Loratadine (Lor), which are histamine receptor 1 (HR1) antagonists that are routinely used for the treatment of human allergy ^57–61^. Eba (and its main metabolite carebastine) and Lor (and its main metabolite desloratadine) can stabilize MCs and block their release of inflammatory mediators ^62–67^. Our results showed that prior-treatment of LAD2 cells with Eba or Lor blocked both Spike-RBD- and Spike-pseudotyped lentivirus- induced degranulation (Figure 6A).

**Figure 6.**
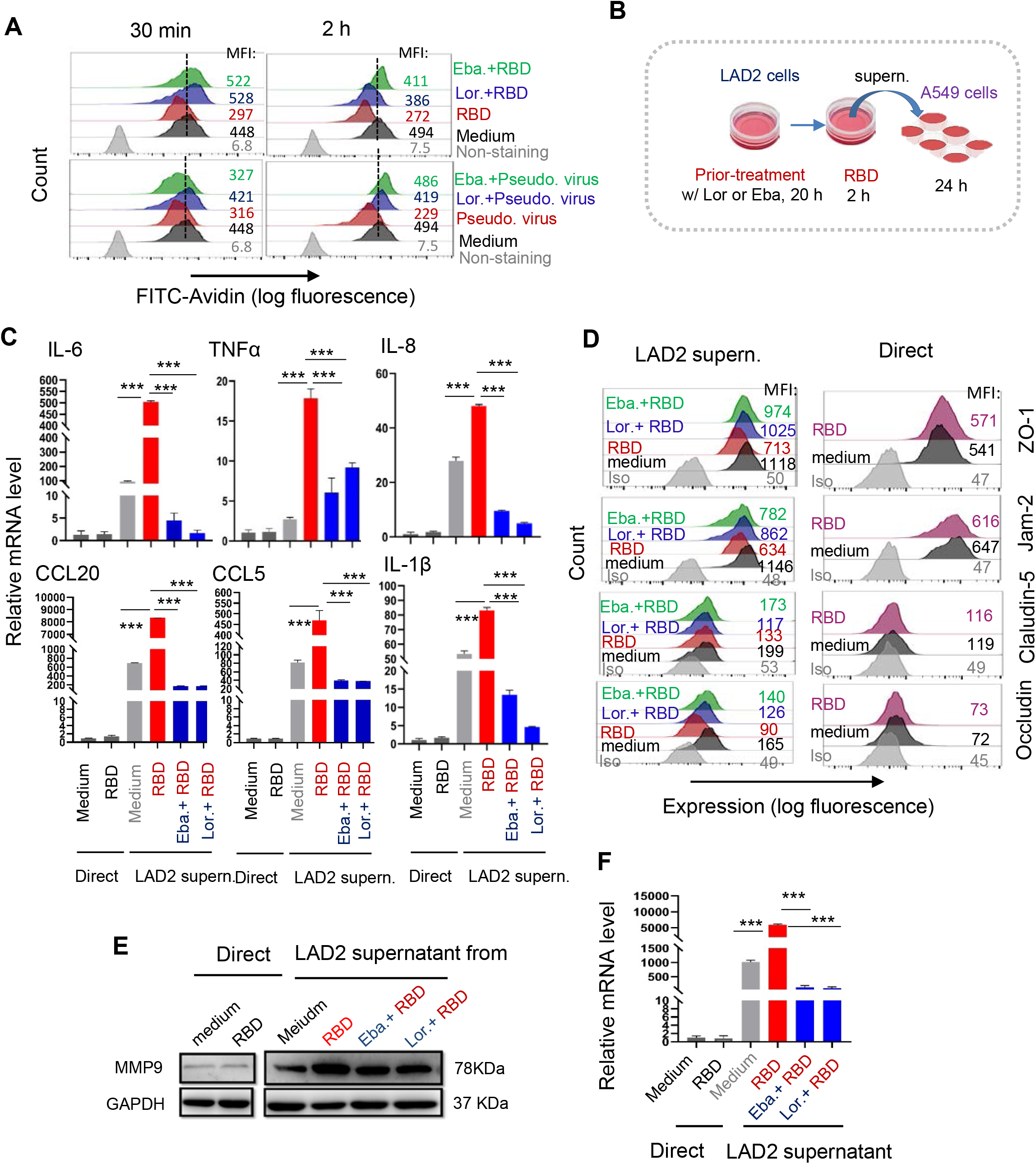
Inhibition of MC degranulation abolishes alveolar epithelial inflammation and protects the tight junction proteins. (**A**) Loratadine and ebastine inhibit Spike-RBD or pseudotyped lentivirus-induced LAD2 cell degranulation. Cells were prior-treated with Loratadine (5 μg/mL) or Ebastine (3 μg/mL) for 20 h, then were incubated with Spike-RBD (5 μg/ml) or Spike-pseudotyped lentivirus (5 ng p24^Gag^) at 37°C for the indicated time, and cell degranulation was detected with Flow cytometry. (**B-F**) The effects of MC degranulation on lung epithelial cells. The illustration for treatments (B). LAD2 cells were prior-treated with or without Loratadine (5 μg/mL) or Ebastine (3 μg/mL) for 20 h, then cells were treated with Spike-RBD (5 μg/ml) for 2 h, and the culture supernatants were harvested to treat A549 cells for additional 24 h, or A549 cells were directly treated with or without Spike-RBD for 24 h. The mRNA levels of IL-6, TNF-α, IL-8, CCL20, CCL5 and IL-1β were quantified with real time q(RT-) PCR, and normalized to *gapdh* mRNA (C). The expressions of ZO-1, Jam-2, Claudin-5 and Occludin were detected by immunstaining with specific antibodies and analyzed with flow cytometry (D). The expression of MMP9 in A549 cells was measured with Western blotting and real time q(RT-) PCR (E, F). Data are presented as mean ± SD. *** p<0.001 is considered significant difference. MFI: mean fluorescence intensity.

From transcriptome analyses, we found linkage between the Spike-RBD-triggered MC degranulation and the alveolar epithelial inflammation and structural impairments. As expected, the use MC stabilizers prevented virus-induced MC degranulation and thus reduced the inflammatory responses and impairment of lung epithelial cells (Figure 6B).

The direct stimulation of A549 cells with Spike-RBD did not induce obvious expression of pro-inflammatory cytokines, however the treatment of A549 cells with the RBD/LAD2 supernatants induced production of extremely high level of IL-6, TNF-α, IL-8, IL-1β, CCL20 and CCL5 (Figure 6C; Figure S3). When LAD2 cells were first treated with Lor (5 μg/mL) or Eba (3 μg/mL) and then subjected to RBD binding, the harvested supernatant lost its capacity to induce pro-inflammatory cytokines due probably to a blockade of degranulaiton (Figure 6C; Figure S3).

The disruption of integrity of alveolar epithelial barrier is associated with defects in the tight junction proteins. In keeping with the transcriptome analysis data, we found that the stimulation of A549 cells with RBD/LAD2 supernatants reduced the tight junction protein ZO-1, Jam-2, Claudin-5 and Occludin, but the direct stimulation with RBD proteins showed no effect; with prior-treatment of LAD2 with Lor or Eba to block degranulation, the harvested RBD/LAD2 culture supernatants were unable to impair tight junction proteins (Figure 6D). These suggest that specific factor or factors released into the RBD/LAD2 supernatants acted on the tight junction proteins.

A possible candidate is Matrix Metallopeptidase 9 (MMP-9) which has significantly elevated expression and activity in COVID-19 patients ^68^. MMP-9 can impair the alveolar epithelial-endothelial capillary barrier by degrading the extracellular matrix, stimulates neutrophil and leukocyte migration, and promote inflammation ^69^. Consistent with this idea, the RBD/LAD2 supernatants- treated A594 cells have high MMP9 expression levels (Figure 6E, 6F), and after blocking LAD2 degranulation with Lor or Eba, the RBD/LAD2 supernatants lost its ability to induce MMP9 gene expression (Figure 6E, 6F).

Taken together, the above data suggest a mechanism whereby RBD-binding to ACE2 molecule induces LAD2 cells degranulation and release of cytokines including MMP9; upon transfer to A549 alveolar epithelial cells, MMP9 disrupts tight junction proteins. In keeping with this mechanistic interpretation, drugs that preventing MC degranulation can reduce alveolar epithelial inflammation and protect the tight junction proteins integrity.

### MC stabilizers reduce SARS-CoV-2-induced lung inflammation and prevent lung injury in mice

To test whether the above observed molecular mechanism derived from *in vitro* study applies to actual infection *in vivo*, we went back to the SARS-CoV-2 infection model of hACE-2 humanized mice. The C57BL/6N-Ace2^em2(hACE2-WPRE,pgk-puro)/CCLA^ were treated with Lor (10 mg/kg) or Eba (5 mg/kg) 1 day prior to intranasal infection with SARS-CoV-2 (strain 107) at a dose of 5×10^6^ TCID_50_, and then the MC stabilizers were continued administered daily for 5 days until the mice were euthanized at 5 dpi (Figure 7A). In untreated controls, SARS-CoV-2 challenge induced MC accumulation in the peri-bronchus, and robust MC degranulation was evidenced by abundant released granules in peri-bronchus and scatted in alveolar spaces (Figure 7B, 7H). The H.E. staining of the adjacent lung sections showed lung lesions including the alveolar septal thickening, inflammatory cell infiltration, fibrinous exudate, hyaline film formation, mucosa desquamation, and hemorrhage (Figure 7C). The administration of Lor and Eba blocked MC degranulation (Figure 7D, 7F), and greatly reduced lung lesions (Figure 7E, 7G, and 7H). Additionally, the administration of these two MC stabilizers significantly dampened SARS-CoV-2-induced inflammation, as evidenced by the greatly reduce production of IL-6, TNF-α, CCL20, CCL5, IL-8, IL-1β, IFN-γ and CRP post viral infection (Figure 7I, Figure S5).

**Figure 7.**
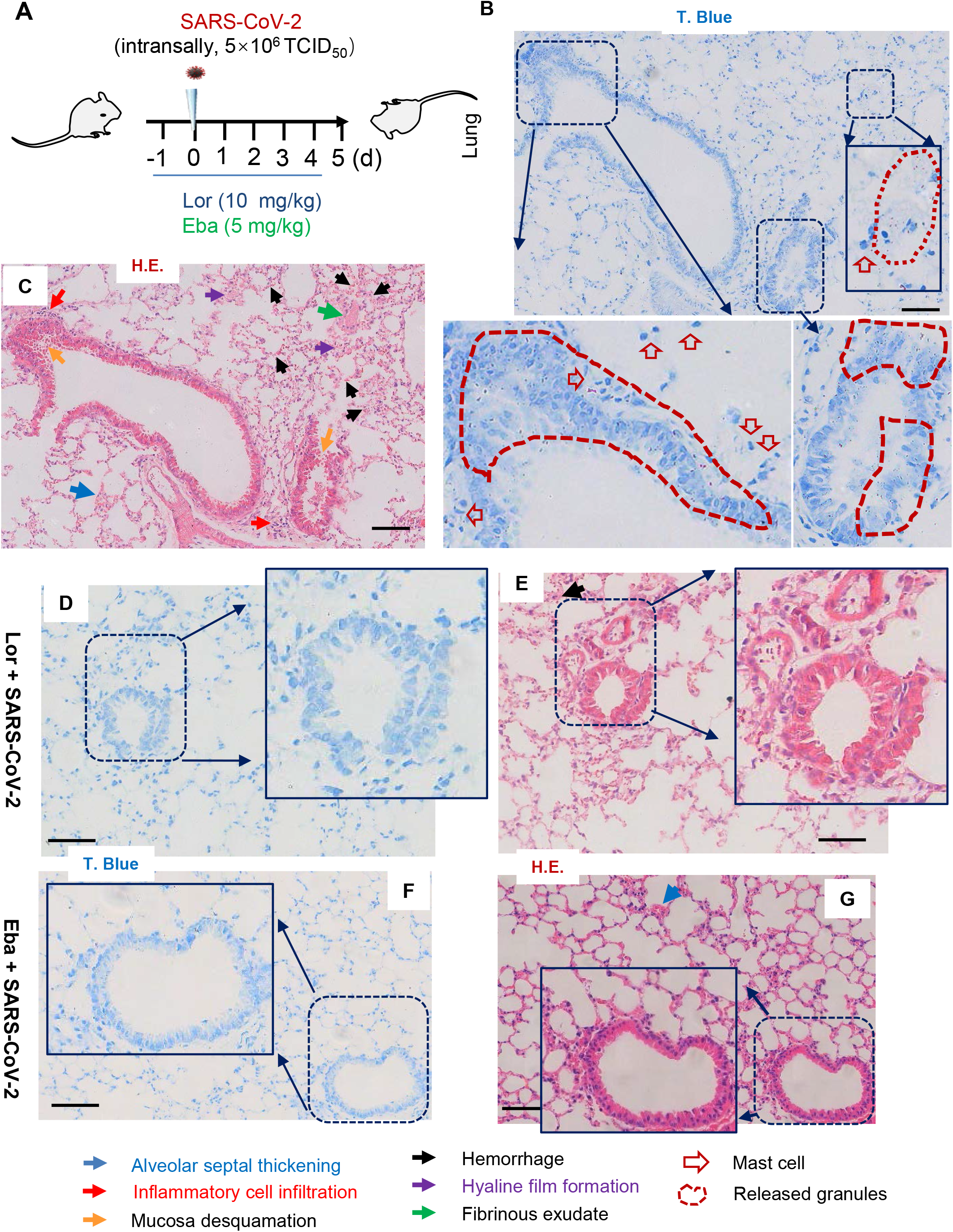
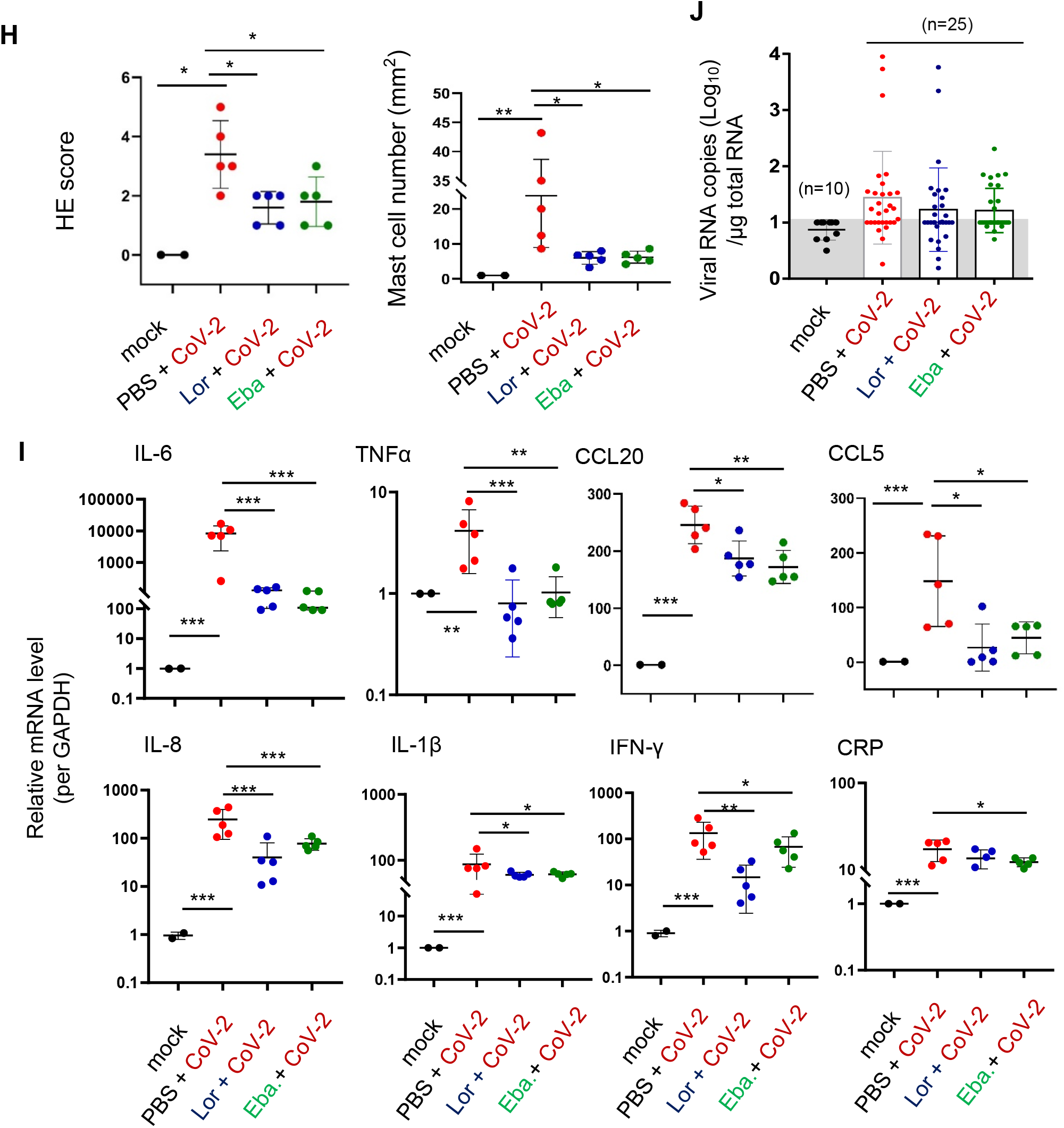
MC stabilizers reduce SARS-CoV-2-induced inflammation and prevent lung injury in mice. (**A**) The illustration of mice treatment. C57BL/6N-Ace2^em2(hACE2-WPRE,pgk-puro)/CCLA^ mice were prior-administered with or without Ebastine (5 mg/kg) or Loratadine (10 mg/kg) 1 day before intranasal infection with SARS-CoV-2 (strain 107) at a dose of 2×10^6^ TCID_50_, and the Ebastine and Loratadine treatments were continued each day over the couse of infection. 5 mice for each treatment groups, and 2 mice without infection and drug treatment were used as the mock controls. Mice were euthanized and lung lobes were harvested for the followed analysis. (**B**-**G**) Pathological lung section. Toluidine blue staining was used to observe MC degranulation, and the lung injury was observed via H.E. staining. Scale bar: 100 μm. (**H**) H.E. scores and MC counts in lung section (5 dpi); (**I**) The mRNA levels of IL-6, TNF-α, CCL20, CCL5, IL-8, IL-1β, IFN-γ, and CRP were quantified with q(RT-) PCR, and normalized to *gapdh* mRNA. (**J**) Viral replication. Total RNAs were prepared from the left lobe, the right lobus superior, the right lobus medius, the right lobus inferior and the pulmonary azygos lobe of each mouse, viral replication was monitored by quantifying the expression of nucleocapsid gene. Data are presented as mean ± SD. * p<0.05, ** p<0.01 and *** p<0.001 are considered significant differences.

Viral replication was also monitored. Total RNAs were prepared from the left lobe, the right lobus superior, the right lobus medius, the right lobus inferior and the pulmonary azygos lobe of each mouse, and viral replication was measured by quantifying the expression of nucleocapsid gene. The mono-treatment with either Lor or Eba had not significantly reduced viral replication (Figure 7J). Consistent with the idea that these drugs reduce SARS-CoV-2-induced lung inflammation and injury directly through preventing MC degranulation and subsequent inflammation, rather than indirectly by reducing virus load.

## Discussion

In this study, we demonstrate a pivotal role of MCs in SARS-CoV-2-induced epithelial inflammation and lung injury in *vivo* and elucidated its possible mechanisms in *vitro*. The validation of MC stabilizers Eba and Lor can inhibit SARS-CoV-2-induced hyper-inflammation and lung injury is highly significant, because it suggests an off-label use of MC stabilizers as immunomodulators to treat the severe cases of COVID-19.

Eba and Lor are the second generation antihistamines that are routinely used clinically for treating allergy, and the higher dosage of these antihistamines can be used as MC stabilizers to block their degranulation ^57,62–67^. Our data in cell lines and in animal models clearly demonstrated a beneficial role of their usage in the setting of SARS-CoV-2 infection, the next step should be clinical testing of these and other MC stabilizers or higher dosage of antihistamines in the treatment of lung injury in COVID-19 patients.

The strategical location at mucosa makes MC being the sentinel and first responders to exposed pathogens. SARS-CoV-2 is not the only pathogen that induce MC degranulation. A variety of pathogens including DNA/RNA virus, fungi, bacteria or their products can activate MCs, and induce secretion of cytokines and chemokines through degranulation-dependent or degranulation-independent pathways ^44^. On the one hand, the released mediators drive the recruitments of effectors cells such as NK cells, multiple CD4^+^ T cell subsets, protective CD8^+^ T cells, for pathogens clearance; on the other hand, these inflammatory mediators can be detrimental as the induced inappropriate inflammatory responses may disrupt the epithelial-endothelia barriers to promote pathogens invasion ^44^. MCs showed cross-talk with multiple types of immune cells via direct cell-to-cell contact or indirectly through the release of soluble factors ^42,43^. Given our findings of SARS-CoV-2 Spike-RBD induced rapid MC degranulation, one may speculate that the MC activation and consequential inflammatory mediators release is one of the root-causes of lung inflammation and injury in COVID-19.

The phenotypes of MC degranulation-induced epithelial inflammation and lung injury in our study is concordant with the immunological and histopathological findings of COVID-19 patients and infected animals ^45,70,71^, indicating that SARS-CoV-2-induced MC degranulation is a major cause that triggers hyper-inflammation and lung injury. We find that MC degranulation induces extremely high level of productions of pro-inflammatory cytokines and chemokines, including IL-6 which has been shown to be an independent risk factor for disease severity and death of COVID-19 ^14,18,21–23^.

Spike-RBD-triggered MC degranulation provides an important route to disrupt alveolar barrier by inducing various types of metallopeptidases. RBD/LAD2 supernatant promotes the expression of MMP9 in alveolar epithelial cells, which resembles the observation of significantly increased circulating MMP9 level in COVID-19 patients ^68,72^. In acute lung injury, the released MMP9 are able to promote degradation of the alveolar-capillary barrier ^69^, through probably action on tight junction proteins.

The released components of SARS-CoV-2-induced MC degranulation contain Chymase and Typtase, which can also be used to explain virus-induced permanent damage to the alveoli epithelial cells and capillary endothelial cells. In inflammation or pathological conditions, the released Chymases from activated MCs are able to amplify local angiotensin-2 concentration to induce inflammatory leucocyte recruitment and endothelial dysfunction, which would further increase pulmonary vascular permeability and cause lung injury ^73,74^. Other types of viruses such as Dengue virus has been reported to trigger the release of Chymase and Typtase to induce breakdown of endothelial cell tight junctions and promote vascular permeability ^75^.

SARS-CoV-2 infection can shutdown the expressions of cell cycle kinases (e. g. CDK1/2/5 and AURKA, *etc*) and result in cell cycle arrest between S and G2 phases of the cell cycle ^70,76^. In our study, we find that Spike-RBD-triggered MC degranulation shutdown the expressions of various genes linking to the regulation of cell cycles/division. Besides, RBD/LAD2 supernatants can directly suppress the expression of genes encoding the cytoskeleton/microtubule-associated proteins and myosins, suggesting MC degranulation induces the change of cytoskeleton organization. Our experimental data, again, consistent with the clinical observation that SARS-CoV-2 infection inducing changes of cytoskeletal-microtubule organization ^70,76^.

It seems that the infection and replication of SARS-CoV-2 in MCs is not necessarily required for triggering degranulation, as the stimulation of Spike-RBD can trigger MC degranulation. MC cells express ACE2 receptor and are permissive for SARS-CoV-2 replication, whether the replication of SARS-CoV-2 in MCs leads to *de novo* synthesis of inflammatory mediators for the secondary release is worthy of further investigation. Additionally, SARS-CoV-2-induced MC degranulation may also be used to explain the disease severity in aged individuals. We find that SARS-CoV-2 induces a more robust MC degranulation in aged chRMs, which could link the severer cytokine storm and higher immune cell infiltration in older adults and in aged RMs ^52,71,77^.

The limitation of our study is that the administration of Eba or Lor significantly reduced SARS-CoV-2-induced inflammation and lung injury, but showed limitation in suppressing viral replication in lungs of ACE2 humanized mice. A combination of MC stabilizers with antiviral drugs such as the RNA polymerase inhibitors Remdesivir and Favipiravir ^6,78–80^, may provide more optimal treatment strategy that dampening inflammation and clearing viruses at the same time.

In summary, we demonstrate that SARS-CoV-2 triggers lung MCs degranulation, which induces the remodeling of various cellular signalings in human alveolar epithelial cells, particularly, MCs degranulation induces the alveolar epithelial inflammation and leads to the consequent disruption of tight junction proteins; importantly, we find that the clinically used MC degranulation stabilizers Eba and Lor are potent agents at reducing virus-induced production of pro-inflammatory factors and preventing lung injury. Our finding uncovers a potentially novel mechanism of SARS-CoV-2 infection initiates alveolar epithelial inflammation and induces lung Injury. Significantly, our results suggest a potential off-label use of MC stabilizers as immunomodulators to treat the severe cases of COVID-19.

## Methods

### Ethics statement

All animal experiments were approved by the Institutional Animal Care and Use Committee of XXXXXX. The SARS-CoV-2 animal model experiments and protocols were also discussed explicitly and extensively with biosafety officers and facility managers. All animal experiments and wild type virus were conducted within the animal biosafety level 3 (ABSL-3) facility.

### Cells and viruses

Human mast cell LAD2 was cultured in RPMI 1640 medium (Gibco) containing 10% fatal bovine serum (FBS) (Gibco) with 100 U/mL penicillin and 100 μg/mL streptomycin. For degranulation experiment, LAD2 cells were grown in StemPro-34 medium (Gibco) supplemented with 100 μg/ml SCF (Novoprotein), 100 μg/ml IL-6 (Novoprotein), nutrient supplement (NS) (Gibco), 100 U/ml penicillin (Invitrogen), 100 μg/ml of streptomycin (Invitrogen) and 2 mM L-Glutamine (Gibco). Adenocarcinomic human alveolar basal epithelial cells (A549) was cultured in DMEM/F12 medium (Gibco) containing 10% FBS (Gibco) with 100U/mL penicillin and 100μg/mL streptomycin. All cells were incubated at 37 °C with 5% CO_2_.

The wild type virus SARS-CoV-2 (2019-nCoV WIV04) was isolated by Wuhan Institutes of Virology, Chinese Academy of Sciences ^53^. The 107 strain of SARS-CoV-2 was provided by Guangdong Provincial Center for Disease Control and Prevention, Guangdong Province of China. Pseudotyped virus was generated by EZ Trans cell transfection reagent-mediated co-transfection of HEK293T cells with the Spike-expressing plasmid pcDNA3.1-2019-nCoV-S-IRES (strain 2019-nCoV WIV04) and pNL4-3. Luc. ΔR ΔE ^81^. These two plasmids are provided by Dr. Lu Lu (Fudan University, Shanghai, China). Harvested supernatants of transfected cells that contained viral particles were aliquoted and stored at −80 °C.

### ACE2-humanzied mice and rhesus macaques experiments

3-4 months old C57BL/6N-Ace2^em2(hACE2-WPRE,pgk-puro)/CCLA^ mice were provided by Guangzhou Institutes of Biomedicine and Health, Chinese Academy of Science 47. Mice (5 for each groups) were infected nasal inhalation with SARS-CoV-2 (strain 107)(5×10^6^ TCID_50_)for indicated times. In some mice, ebastine (5 mg/kg) or loratadine (10 mg/kg) (both from Sigma-Aldrich) was administered 1 day before infection and the treatments were continued each day over the course of infection. The lungs were collected on the day of euthanization for pathological, virological, and immunological analysis.

For Ad5-hACE2-transduced BALB/c mice ^51^, specific pathogen-free 6-10 weeks old male and female BALB/c mice were lightly anesthetized with isoflurane and transduced intranasally with 2.5×10^8^ fluorescence focus units (FFU) of Ad5-ACE2 in 75 μL DMEM. Five days post-transduction, mice were infected intranasally with 7×10^4^ TCID_50_ SARS-CoV-2 in a total volume of 50 μL DMEM. The SARS-CoV-2 strains used in this experiment were isolated from COVID-19 patients in Guangzhou and in Washington state (Accession numbers: MT123290, MN985325.1). At the indicated time, the lungs of mice were collected on the day of euthanization for histology analysis.

For monkey study, eight Chinese rhesus macaques (chRMs) (*Macaca mulatta*), including young group (3- to 6-year old) and aged group (17- to 19-year old), were anaesthetized by Zoletil 50 (Viabac, France) and intratracheally inoculated with SARS-CoV-2 (virus stain107) (1□×□10^7^ TCID_50_) in a 2 mL volume by bronchoscope. The animals were euthanized at 7 or 15 dpi and the lung lobes were collected for histology analysis ^52^.

### Histology

Animal’s lungs were fixed in zinc formalin. For routine histology, tissue sections (~4 μm each) were stained with Hematoxylin or Eosin or Toluidine blue. The pathological score was assessed according the degree of lung tissue lesions including alveolar septal thickening, hemorrhage, inflammatory cells infiltration, and consolidation. The semiquantitative assessment were performed as follows^47^, 0: no alveolar septal thickening; 1: alveolar septal thickening was very mild, the area of alveolar septal thickening, hemorrhage and inflammatory cells infiltration was less than 10%; 2: when alveolar septal thickening was mild, the area of alveolar septal thickening, hemorrhage and inflammatory cells infiltration was 10%-25%; 3: when alveolar septal thickening was moderate, the area of alveolar septal thickening, hemorrhage, inflammatory cells infiltration, hyaline film formation, mucosa desquamation and fibrinous exudate was 25%-50%; 4: when alveolar septal thickening was marked, the area of alveolar septal thickening, hemorrhage, inflammatory cells infiltration, hyaline film formation, mucosa desquamation and fibrinous exudate was 50%-75%; 5: when alveolar septal thickening was very marked, the area of alveolar septal thickening, hemorrhage, inflammatory cells infiltration, hyaline film formation, mucosa desquamation and fibrinous exudate was greater than 75%.

### Spike-RBD protein binds to LAD2 cells

LAD2 cells (3×10^5^) were incubated with Spike-RBD protein (5 μg/mL, Genscript, Z03483) in adherent buffer (1mM CaCl2, 2mM MgCl_2_ and 5% BSA, pH 7.4) for 1h at 4 °C. The cells were then fixed with 4% paraformaldehyde (Sigma-Aldrich) for 30 min at room temperature and stained with anti-His-tag antibodies (Abmart, M30111S). Subsequently, the cells were stained with goat anti-mouse Alexla Fluor 488-conjugated secondary antibodies (Invitrogen, A11001), and were detected with flow cytometry. In some experiments, LAD2 cells were prior-treated with 0.25% trypsin (without EDTA) for 10 min at 37 °C or prior-blocked with anti-ACE2 antibody (5 μg/mL, R&D Systems, AF933) for 2 h at 37 °C before the incubation with Spike-RBD protein.

### LAD2 cell degranulation

LAD2 cells (3×10^5^) were exposed to Spike-RBD protein (Genscript) (5 μg/mL), nucleocapsid protein (provided by Jincun Zhao, Guangzhou Medical University ^51^ (5 μg/mL), or SARS-CoV-2(M.O.I.=1) (strain 2019-nCoV WIV04) ^53^, for the indicated times. Mast cell degranulation activator compound 48/80 (C48/80) (4 μg/ml) (Sigma, C2313) was used as the control. Cells were fixed with 4% paraformaldehyde (Sigma-Aldrich) at room temperature for 30 min, and washed 3 times with PBS, then cells were incubated with anti-avidin-FITC (500 ng/mL, Invitrogen, A821) which was diluted in permeabilized buffer (1% FBS and 0.2% Triton X-100 in PBS) at 4 °C for 1 h. After washing, cells were detected with BD Accuri C6 and analyzed with FlowJo. In some experiments, Loratadine (5 μg/mL, Selleck) or Ebastine (3 μg/mL, Selleck) was used to prior-treat cells for 20 h before stimulation with Spike-RBD protein. The LAD2 cell culture supernatants were harvested for quantifying the released components of Chymase and Tryptase with ELISA kits according the manufacturer`s instructions instruments (Lunchangshuo Biotech, Tryptase: SU-B10563; Chymase: SU-B16617).

### Inflammatory cytokine assay

A549 cells (3×10^5^) were treated with LAD2 culture supernatant (250 μL) for 24 h, then cells were harvested. The cytokines were determined either by quantifying the production of mRNAs or intracellular immuno-staining with specific antibodies. For immuno-staining assay, A549 cells were added leukocyte activation cocktail containing BD GolgiPlug (BD, 550583) and cultured for 6 h. Following activation, cells were washed with FACS buffer. BD Cytofix/Cytoperm solution (BD, 554722) was used for the simultaneous fixation and permeabilization of cells for 20 min at 4°C before intracellular cytokine staining. Antibodies diluted in Perm/Wash buffer was added and cells were further incubated at 4°C for overnight. After washing, cells were resuspended in FACS buffer to flow cytometric analysis. Cytokine antibodies against the flowing markers were used: Alexa Fluor 647-IL-1β (Biolegend, JK1B-1), PE-IL-6 (BD, MQ2-6A3), BV421-IL-8 (BD, G265-8).

### Real time (RT-) PCR

Total RNAs were extracted by using TRIzol Reagent (Invitrogen) and then reverse transcribed into cDNA with synthesis Kit (TOYOBO, FSQ-301), according to the manufacturer`s instructions. Real-time PCR was carried out by using the SYBR qPCR Mix (Genestar, A33-101) with the following thermal cycling conditions: initial denaturation at 95°C for 2 min, amplification with 40 cycles of denaturation at 95°C for 15s, primer annealing at 60°C for 15 s, and extension at 72°C for 30s. The data were analyzed by green-based SYBR, semi-quantified and normalized with GAPDH. Real-time PCR was performed on the Bio-Rad CFX96 Real-Time PCR system.

The human primers used: IL-6-F, 5′-CAG ACA GCC ACT CAC CTC TTC AG-3′; IL-6-R, 5′-CAG CCA TCT TTG GAA GGT TCA G-3′; TNF-α-F, 5′-CCC AGG CAG TCA GAT CAT CTT C-3′; TNF-α-R, 5′-GTG AGG AGC ACA TGG GTG GAG-3′; MMP9-F, 5′-CCT TCT ACG GCC ACT ACT GTG C-3′; MMP9-R, 5′-GCC AGT ACT TCC CAT CCT TGA AC-3′; GAPDH-F, 5′-ATC CCA TCA CCA TCT TCC AGG-3′; GAPDH-R, 5′-CCT TCT CCA TGG TGG TGA AGA C-3′; IL-1β-F, 5′-CGT CAG TTG TTG TGG CCA TGG A-3′; IL-1β-R, 5′-GAG CGT GCA GTT CAG TGA TCG TA-3′; IL-8-F, 5′-CTG ATT TCT GCA GCT CTG TGT GA-3′; IL-8-R, 5′-GGT CCA GAC AGA GCT CTC TTC CA-3′; CCL20-F, 5′-GCT GCT TTG ATG TCA GTG CT-3′; CCL20-R, 5′-TGT CAC AGC CTT CAT TGG C-3′; CCL5-F, 5′-ACC ACA CCC TGC TGC TTT G-3′; CCL5-R, 5′-GCG GTT CTT TCG GGT GAC A-3′.

The mouse primers used: IL-6-F, 5′-CTT CCA TCC AGT TGC CTT CTT G-3′; IL-6-R, 5′-AAT TAA GCC TCC GAC TTG TGA AG-3′; TNF-α-F, 5′-CAG ACC CTC ACA CTC AGA TCA TCT-3′; TNF-α-R, 5′-CCT CCA CTT GGT GGT TTG CTA-3′; IL-1β-F, 5′-CTT TCA GAG GCC AGA GAG TCC-3′; IL-1β-R, 5′-TCC CTG TAG TGA CAG CAC CT3′; IL-8-F, 5′-CGG CAA TGA AGC TTC TGT AT-3′; IL-8-R, 5′-CCT TGA AAC TCT TTG CCT CA-3′; INFγ-F, 5′-ATG AAC GCT ACA CAC TGC ATC-3′, INFγ-R, 5′-CCA TCC TTT TGC CAG TTC CTC-3′; GAPDH-F, 5′-TGC ACC ACC AAC TGC TTA G-3′; GAPDH-R, 5′-GAT GCA GGG ATG ATG TTC-3′; CRP-F, 5′-CAG AGA TTC CTG AGG CTC CAA CA-3′; CRP-R, 5′-AGT CAC CGC CAT ACG AGT CCT G-3′; CCL20-F, 5′-AAG ACA GAT GGC CGA TGA AG-3′; CCL20-R, 5′-AGG TTC ACA GCC CTT TTC AC-3′; CCL5-F, 5′-GTG CCC ACG TCA AGG AGT AT-3′; CCL5-R, 5′-GGG AAG CTA TAC AGG GTC A-3′.

The mice extracted RNAs were used to measure the copies of nucleocapsid gene SARS-CoV-2 using THUNDERBIRD Probe One-step qRT-PCR Kit (Toyobo).The primer sequences for qRT-PCR were as follows: Forward primer: 5′-GGG GAA CTT CTC CTG CTA GAA T-3′; Reverse primer: 5′-CAG ACA TTT TGC TCT CAA GCT G-3′. The TaqMan probe sequences were 5′-FAM-TTG CTG CTG CTT GAC AGA TT-TAMRA-3′. The standard samples were purchased from the National Institute of Metrology of China.

### Western blotting

LAD2 or A549 cells were lysed for 1 h at 4 °C in lysis buffer (Beyotime). After centrifugation for 10 min at 12, 000 g, the supernatant was boiled in reducing SDS sample loading buffer and analyzed by SDS-PAGE. The anti-ACE2 antibody (Abcam, EPR4435), anti-MMP9 antibody (signal antibody, 29091), anti-GAPDH antibody (Abcam, 6C5), and the horseradish peroxidase-conjugated secondary antibody were used in Western blotting.

### Flow Cytometry

The expression of ACE2 in LAD2 was determined by immunostaining with PE-labeled rabbit anti-ACE2 (Bioss, bs-1004R) and detecting with flow cytometry. For detecting tight junction proteins ZO-1, Occludin, Claudin-5 and JAM2 in A549 cells, cells were blocked with 5% BSA in PBS for 1 h at room temperature then incubated with primary antibodies for 2h at 4°C. Primary antibodies against ZO-1 (Invitrogen, 402200), Occludin (Invitrogen, OC-3F10), Claudin-5 (Invitrogen, 4C3C2) and JAM-2 (Abcam, EPR2489), were used. A permeabilizing agent (1%FBS and 0.2% Triton X-100 in PBS) was used for ZO-1 intracellular staining. Cells were washed with FACS buffer and then incubated with Alexa Flour 488-labeled goat anti-rabbit or goat anti-mouse IgG (Invitrogen, A11034; Invitrogen, A11001) for 1 h at 4 °C, then cells were analyzed with flow cytometry.

### RNA Sequencing and data analysis

A549 cells were treated with LAD2 cell culture supernatants for 24 h. Total RNAs were extracted using Trizol (Invitrogen) according to the manufacturer’s protocol, and ribosomal RNA removed using QIAseq FastSelect-rRNA HMR Kits (QIAGEN, Germany). Fragmented RNAs (average length approximately 200 bp) were subjected to first strand and second strand cDNA synthesis, followed by adaptor ligation and enrichment with a low-cycle according to the instructions of NEBNext UltraTM RNA Library Prep Kit for Illumina (NEB, USA). The purified library products were evaluated using the Agilent 2200 TapeStation and Qubit2.0 (Life Technologies, USA). The libraries were paired-end sequenced (PE150, Sequencing reads were 150 bp) at Guangzhou RiboBio Co., Ltd. (Guangzhou, China) using Illumina HiSeq 3000 platform.

Raw RNA sequencing (RNA-seq) reads were filtered using Trimmomatic v0.36 ^82^. The filtered reads were mapped to the human (hg38) reference genomes using HISAT v2.1 with corresponding gene annotations (GRCh38.p13) with default settings ^83^. Total counts per mapped gene were determined using *featureCounts* function in SubReads package v1.5.3 with default parameter ^84^. Next, counts matrix obtained from *featureCounts* was used as input for differential expression gene analysis with the bioconductor package DESeq2 v1.26 in R v4.0^85^. Gene counts more than 5 reads in a single sample or more than 50 total reads across all samples were retained for further analysis. Filtered counts matrix was normalized using the DESeq2 method to remove the library-specific artifacts. Principal component analysis was based on global transcriptome data using the build-in function *prcomp* in R software. The genes with log2 fold change >1 or < −1 and adjusted p value <0.05 corrected for multiple testing using the Benjamini-Hochberg method were considered significant. Transcription-factor enrichment analysis and functional enrichment analysis was performed using Metascape server tool ^86^ (https://metascape.org/gp/index.html#/main/step1). Gene set enrichment analysis (GSEA) were performed using the R package clusterProfiler v3.18.1 ^87^. Protein-protein interaction (PPI) networks of DEGs were built using STRING v11 with a confidence score threshold of 0.7 and visualized with Cytoscape v3.8.1 ^88,89^.

## Data and code availability

All raw RNA-seq data used in this study have been deposited at the National Genomics Data Center (https://bigd.big.ac.cn/bioproject/browse/PRJCA005128).

## Statistical analysis

Graphpad Prism 8.0 (GraphPad Software) was used for statistical analysis. Unpaired (or paired) two-tailed t test was performed to analyze significant differences. Significance levels are indicated as * p<0.05, ** p<0.01, *** p<0.001.

## Author Contribution

## Declaration of interests

The authors have declared that no conflict of interest exists

## Supplemental Materials

Figure S1-S5

## Notes

### Competing Interest Statement

The authors have declared no competing interest.

