## Supplemental information for "SARS-CoV-2-Triggered Mast Cell Rapid Degranulation Induces Alveolar Epithelial Inflammation and Lung Injury"

**Figure legends**

**Figure S1. Virus distribution in SARS-CoV-2-infected humanized mice.** C57BL/6N-Ace2$^{em2(hACE2\text{-}WPRE,pgk\text{-}puro)/CCLA}$ mice were intranasally infected with SARS-CoV-2 (strain 107) at a dose of 2 $\times 10^6$ $TCID_{50}$, and were euthanized at the 1 dpi and 3 dpi. Lung sections were stained with anti-nucleocapsid antibody (red) and DAPI (blue).

**Figure S2. SARS-CoV-2 infection of LAD2 cells.** LAD2 cells were infected with SARS-CoV-2 (M.O.I. =1) for the indiated time, and the cellular levels of nucleocapsid mRNA were quantified with q(RT-)PCR. Data are normalized to β-actin mRNA and expressed as –ΔCt values.

**Figure S3. Cytokine expressions in LAD2 cells.** LAD2 cells were treated with or without Spike-RBD (5 μg/ml) at 37°C for the indicated time, and the mRNA levels of IL-6, IL-8, IL-1β and TNF-α were quantified with q(RT-) PCR, and normalized to *gapdh* mRNA. One representative result from 3 repeats is shown. Data are presented as mean ± SD. * $p<0.05$, ** $p<0.01$ and *** $p<0.001$ are considered significant differences.

**Figure S4. Assay for LAD2 cell degranulation treated by nucleocapsid**

**protein.** LAD2 cells were treated with or without nucleocapsid protein of SARS-CoV-2 (5 μg/ml) at 37°C for the indicated time, then cells were permeabilized and immunostained with anti-avidin-FITC at 4°C for 1 h, and analyzed with flow cytometry. One representative result from 3 repeats is shown. MFI: mean fluorescence intensity.

**Figure S5. Cytokine productions in A549 cells.** LAD2 cells were prior-treated with or without Loratadine (5 μg/mL) or Ebastine (3 μg/mL) for 20 h, then cells were treated with Spike-RBD (5 μg/ml) for 2 h, and the culture supernatants were harvested to treat A549 cells for additional 24 h, or A549 cells were directly treated with or without Spike-RBD for 24 h. The productions of IL-6, IL-8, and IL-1β were measured with intracellular immunostaining with specific antibodies. One representative result from 3 repeats is shown. MFI: mean fluorescence intensity.

### Figure S1

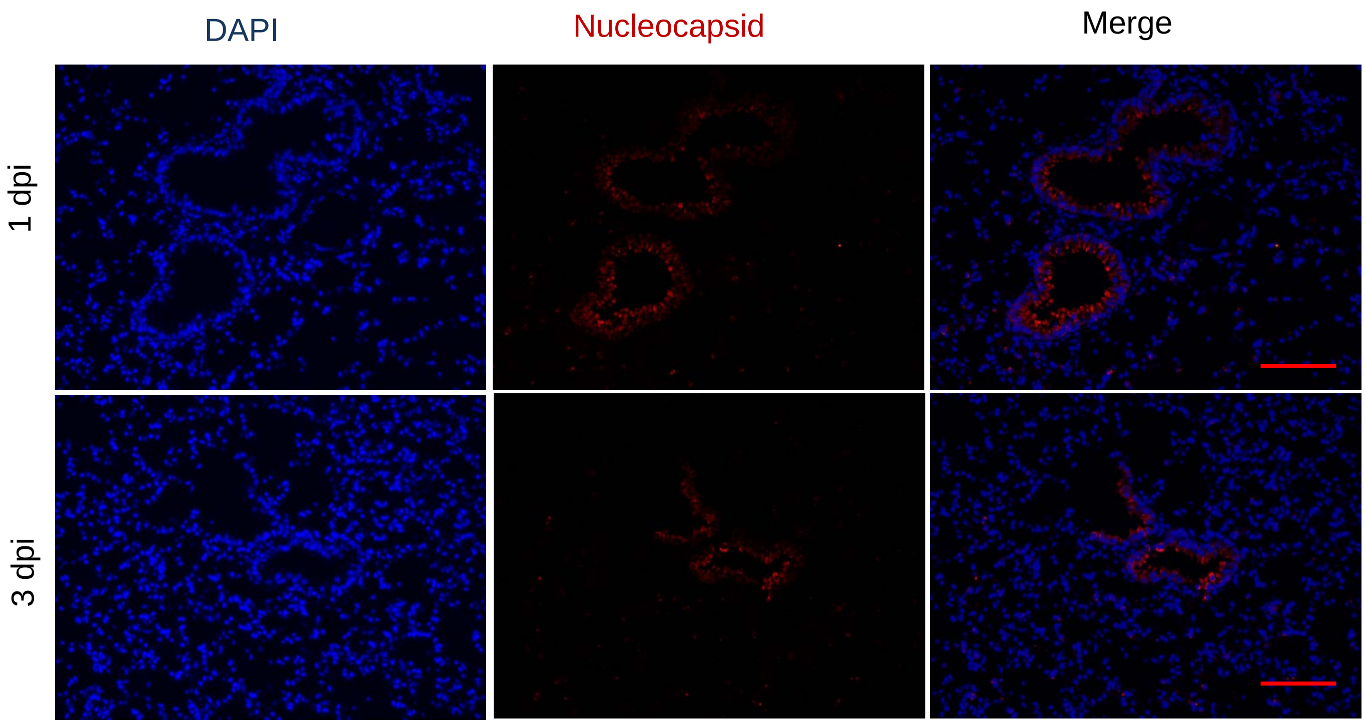

**Figure S2**

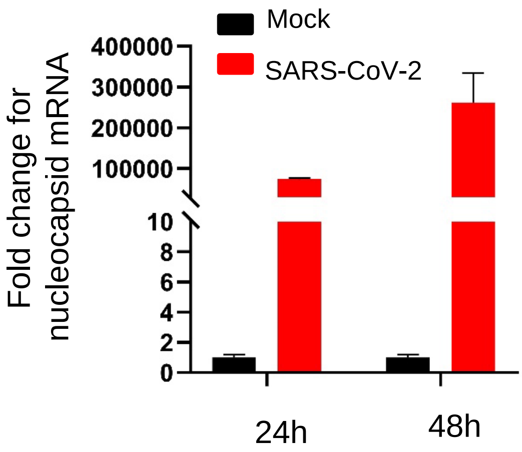
Mock
SARS-CoV-2
Fold change for nucleocapsid mRNA
400000
300000
200000
100000
10
8
6
4
2
0
24h
48h

**Figure S3**

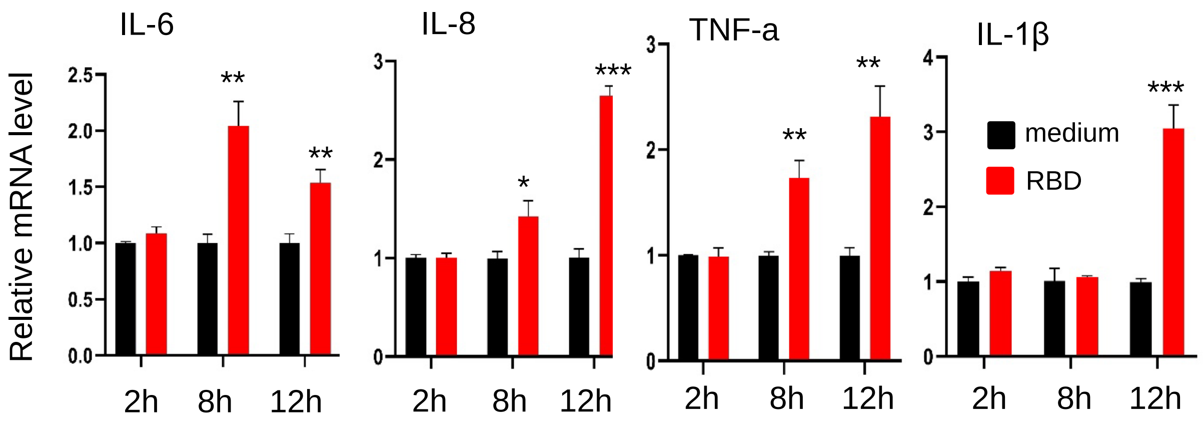
IL-6
IL-8
TNF-a
IL-1β
Relative mRNA level
2.5
2.0
1.5
1.0
0.5
0.0
3
2
1
0
4
**
***
*
medium
RBD
2h
8h
12h

**Figure S4**

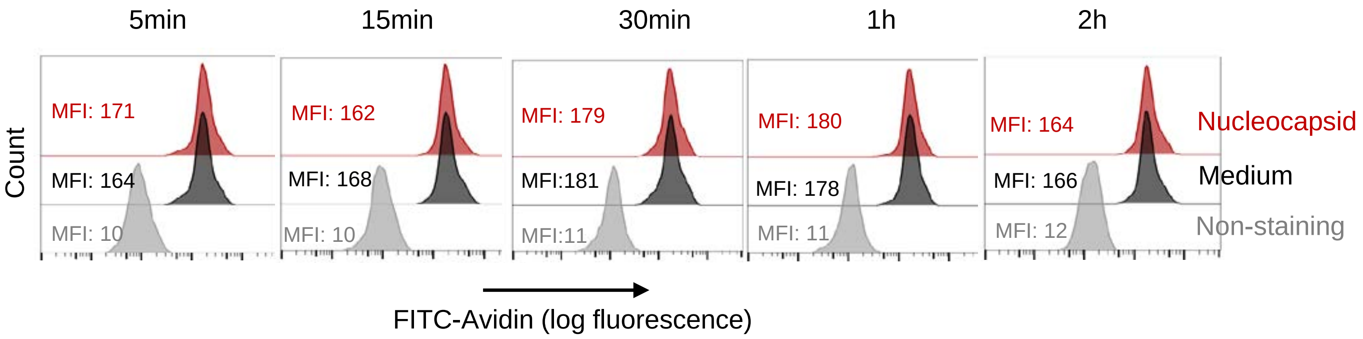
5min
15min
30min
1h
2h
Count
MFI: 171
MFI: 164
MFI: 10
MFI: 162
MFI: 168
MFI: 10
MFI: 179
MFI:181
MFI:11
MFI: 180
MFI: 178
MFI: 11
MFI: 164
MFI: 166
MFI: 12
Nucleocapsid
Medium
Non-staining
FITC-Avidin (log fluorescence)

**Figure S5**

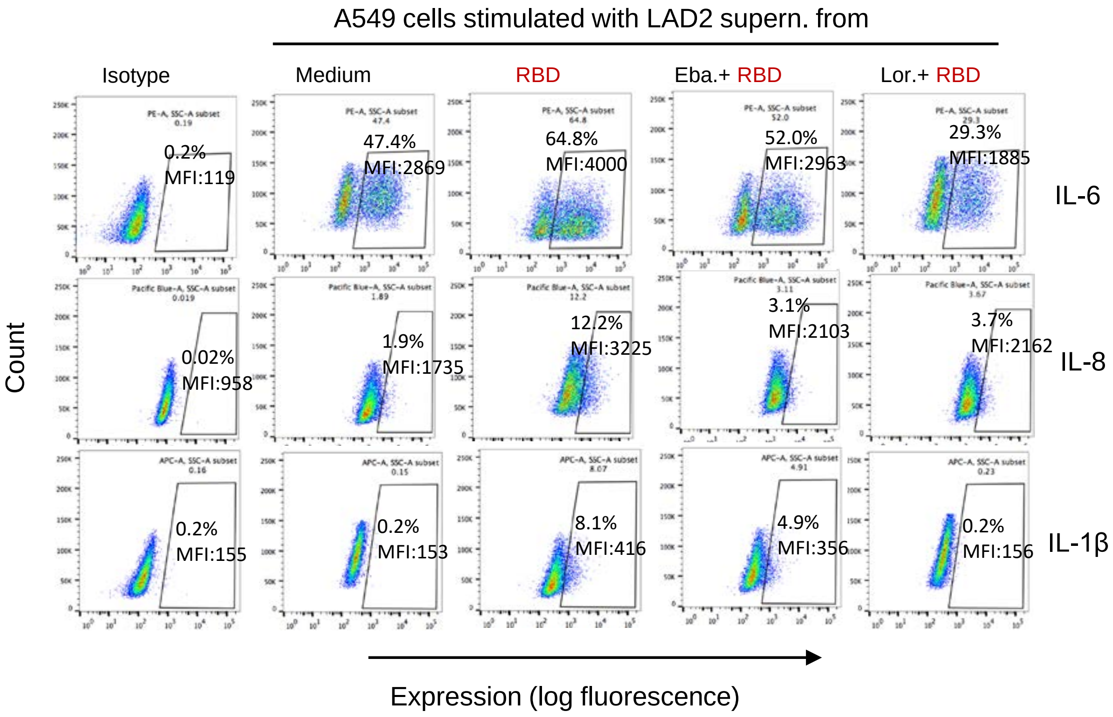
A549 cells stimulated with LAD2 supern. from
Isotype
Medium
RBD
Eba.+ RBD
Lor.+ RBD
0.2%
MFI:119
47.4%
MFI:2869
64.8%
MFI:4000
52.0%
MFI:2963
29.3%
MFI:1885
IL-6
0.02%
MFI:958
1.9%
MFI:1735
12.2%
MFI:3225
3.1%
MFI:2103
3.7%
MFI:2162
IL-8
0.2%
MFI:155
0.2%
MFI:153
8.1%
MFI:416
4.9%
MFI:356
0.2%
MFI:156
IL-1β
Count
Expression (log fluorescence)
